# Multifactorial role of mitochondria in echinocandin tolerance revealed by transcriptome analysis of drug-tolerant cells

**DOI:** 10.1101/2021.07.07.451565

**Authors:** Rocio Garcia-Rubio, Cristina Jimenez-Ortigosa, Lucius DeGregorio, Christopher Quinteros, Erika Shor, David S Perlin

**Affiliations:** Center for Discovery and Innovation, Hackensack Meridian Health, Nutley, NJ, USA; Department of Medical Sciences, Hackensack Meridian Health School of Medicine, Nutley, NJ, USA; Lombardi Comprehensive Cancer Center, Department of Microbiology and Immunology, Georgetown University, Washington DC, USA

**Keywords:** *Candida glabrata*, echinocandins, transcriptomics, mitochondria, antifungal drug tolerance

## Abstract

Fungal infections cause significant mortality and morbidity worldwide, and the limited existing antifungal reservoir is further weakened by the emergence of strains resistant to echinocandins, a first line of antifungal therapy. *Candida glabrata* is an opportunistic fungal pathogen that rapidly develops mutations in the echinocandin drug target, β-1,3-glucan synthase (GS), which are associated with drug resistance and clinical failure. Although echinocandins are considered fungicidal in *Candida*, a subset of *C. glabrata* cells survive echinocandin exposure, forming a drug-tolerant cell reservoir, from which resistant mutations are thought to emerge. Despite their importance, the physiology of rare drug-tolerant cells is poorly understood. We used fluorescence-activated cell sorting to enrich for echinocandin-tolerant cells, followed by modified single cell RNA sequencing to examine their transcriptional landscape. This analysis identified a transcriptional signature distinct from the stereotypical yeast environmental stress response and characterized by upregulation of pathways involved in chromosome structure and DNA topology and downregulation of oxidative stress responses, the latter of which was observed despite increased levels of reactive oxygen species. Further analyses implicated mitochondria in echinocandin tolerance, wherein inhibitors of mitochondrial complexes I and IV reduced echinocandin-mediated cell killing, but mutants lacking various mitochondrial components all showed an echinocandin hypo-tolerant phenotype. Finally, GS enzyme complexes purified from mitochondrial mutants exhibited normal *in vitro* inhibition kinetics, indicating that mitochondrial defects influence cell survival downstream of the drug-target interaction. Together, these results provide new insights into the *C. glabrata* response to echinocandins and reveal a multifactorial role of mitochondria in echinocandin tolerance.

**Importance:** Echinocandin drugs are a first line therapy to treat invasive candidiasis, which is a major source of morbidity and mortality worldwide. The opportunistic fungal pathogen *Candida glabrata* is a prominent bloodstream fungal pathogen and it is notable for rapidly developing echinocandin-resistant strains associated with clinical failure. Echinocandin resistance is thought to emerge within a small echinocandin-tolerant subset of *C. glabrata* cells that are not killed by drug exposure, but mechanisms underlying echinocandin tolerance are still unknown. Here we describe the unique transcriptional signature of echinocandin-tolerant cells and the results of follow-up analyses, which reveal a multifactorial role of mitochondria in *C. glabrata* echinocandin tolerance. In particular, although chemical inhibition of respiratory chain enzymes increased echinocandin tolerance, deletion of multiple mitochondrial components made *C. glabrata* cells hypo-tolerant to echinocandins. Together, these results provide new insights into the *C. glabrata* response to echinocandins and reveal the involvement of mitochondria in echinocandin tolerance.

## INTRODUCTION

Invasive candidiasis is an emerging life-threatening infection recognized as a major cause of morbidity and mortality (1). For decades, antifungal drugs of the azole class have been used as the primary therapy to treat *Candida* infections. Azoles have excellent activity against the predominant *Candida* species, *C. albicans*, and azole resistance in this species remains relatively low. However, an epidemiological shift has been taking place worldwide toward non-*albicans* species that are inherently resistant or readily acquire azole resistance, most notably *Candida glabrata* (2). This shift has led to the widespread use of echinocandin drugs as a first line antifungal therapy to treat and prevent invasive candidiasis (3). Alarmingly, an increase in echinocandin-resistant *C. glabrata* isolates associated with clinical failure has been reported worldwide (4–7). However, the mechanisms enabling *C. glabrata* to develop echinocandin resistance are still very poorly understood.

Although echinocandins are fungicidal drugs in yeast, a small subset of *C. glabrata* cells within a given population demonstrate drug tolerance by surviving prolonged exposures to echinocandins (8, 9). Tolerance to fungicidal drugs is defined as the ability of cells to survive a drug concentration that is expected to kill cells, usually at or above the MIC (8), whereas resistance is manifested as the ability of the cells to grow in the presence of drug and is genetically stable. Clinical resistance to echinocandins is virtually always due to mutations in the hotspot regions of *FKS1* and *FKS2* genes (10). These genetic mutations leading to stable drug resistance are thought to emerge from the drug-tolerant cellular reservoir, making tolerance as a key prerequisite to echinocandin resistance (8, 11). However, the molecular underpinnings of tolerance are currently not understood. Because echinocandins target β-1,3-glucan synthase, an enzyme that helps build the fungal cell wall, thus far studies of echinocandin tolerance have focused on the role of cell wall integrity pathways (12–14). A more unbiased examination of echinocandin tolerance can be derived from exploring the transcriptional landscape of echinocandin-tolerant cells. However, this approach has not been attempted thus far due to the difficulty of studying a very small subset of cells. Instead, the few studies that have examined the transcriptional response to echinocandins have used bulk-RNA isolated from entire echinocandin-treated cultures (15–16), which are largely comprised of dead and dying cells due to the drugs’ quick fungicidal action (17), thus potentially obscuring the transcriptional state of drug-tolerant cells.

In this study, we used fluorescence-activated cell sorting to highly enrich for *C. glabrata* cells that have survived prolonged echinocandin exposure *in vitro*, followed by their transcriptional analysis using a modified single cell RNA sequencing approach wherein transcriptomes of groups of several surviving cells were sequenced. This analysis identified several unique features of the transcriptional landscape of echinocandin-tolerant cells, including a downregulation of the oxidative stress response pathway, which occurs despite an echinocandin-induced increase in reactive oxygen species (ROS) abundance. We also found that, although inhibitors of mitochondrial complexes I and IV reduced both echinocandin-induced ROS formation and echinocandin-mediated cell killing, an ROS scavenger did not alter cell killing dynamics, indicating that ROS per se do not significantly contribute to cell death upon echinocandin exposure. Moreover, we found that mutants lacking various mitochondrial components, including both respiration-proficient and respiration-deficient strains, all showed an echinocandin hypo-tolerant phenotype at sub-MIC concentrations. Finally, we partially purified the echinocandin target enzyme β-1,3-glucan synthase from mitochondrial mutants and found that the enzyme’s sensitivity to echinocandins was not altered, indicating that mitochondrial status influences cell survival downstream of the echinocandin-enzyme interaction. Together, these results provide new insights into the *C. glabrata* response to echinocandins and reveal the involvement of mitochondria in echinocandin tolerance.

## RESULTS

### Using fluorescence-activated cell sorting to enrich for echinocandin-tolerant *C. glabrata* cells for RNA sequencing (RNAseq)

To isolate ultra-rare echinocandin-tolerant *C. glabrata* cells for subsequent transcriptome analysis, we used fluorescence-activated cell sorting (FACS). We tested several combinations of live/dead dyes and fluorescent markers (Table S1) in cells that had been cultured in the presence of either 0.25 µg/ml caspofungin or 0.06 µg/ml micafungin (corresponding to 2X-MIC for strain ATCC2001) for 24 hours, followed by plating of the sorted fractions on drug-free medium and counting the colony forming units. We found that sorting cells that did not stain with propidium iodide (PI) (Figure 1A) resulted in the strongest enrichment for cells capable of producing colonies, from less than one in 1000 events to one in 20-25 events. Thus, the PI-negative cell subset, which contained <1% of all cells, was designated as containing echinocandin-tolerant cells. Next, the sorted cells were analyzed using a modified single cell RNA sequencing approach as follows. Based on pilot experiments comprised of FACS followed by plating and CFU counts, we calculated the number of total events that had to be sorted to obtain approximately ten viable *C. glabrata* cells per well (in order to ensure a robust level of RNA for sequencing). This was done both for echinocandin treated cell samples as well as for untreated controls. Thus, for untreated cells, the wells would contain the same number of viable cells as for echinocandin-treated cells. Although for the latter the wells would also contain some cell debris and inviable cells, viable cells would be significantly enriched in these wells relative to “bulk” RNAseq (see below). Because the echinocandin-tolerant cells were on the one hand expected to be non-growing but on the other hand present in a rich nutritional environment, we used two kinds of “no drug” controls: cells in either logarithmic (mimicking the nutrient-rich environment) or stationary phase (mimicking the non-growing state), stained with PI and sorted in the same fashion (Figure 1A).

**Figure 1.**
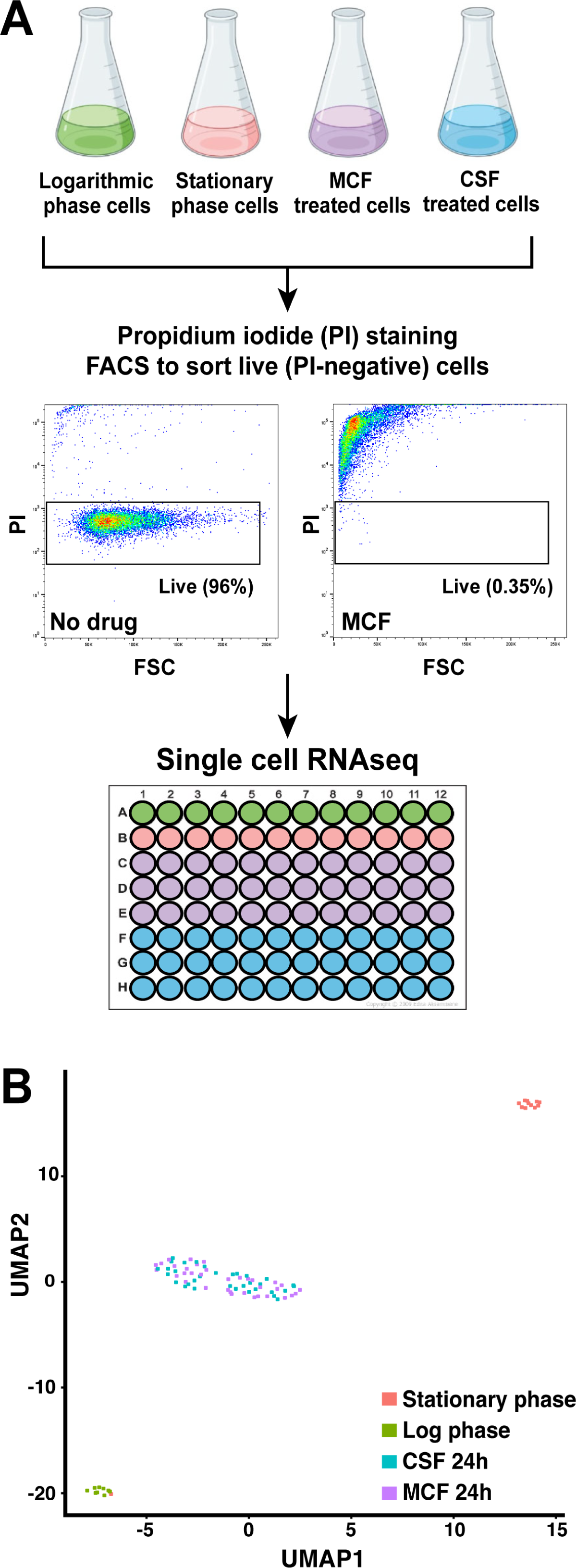
Isolation of rare echinocandin-tolerant *C. glabrata* cells after micafungin or caspofungin exposure followed by single cell RNA sequencing. (A) *C. glabrata* cells were exposed to above-MIC concentrations of caspofungin (0.25 µg/ml) and micafungin (0.06 µg/ml) for 24 hours. Propidium iodide (PI)-negative cells, which accounted for <1% of all cells, were defined as echinocandin-tolerant. Two kinds of “no drug” controls were included: cells in either logarithmic or stationary phase. Sorted cells were analyzed using a modified single-cell RNA sequencing approach where each well contained 5-10 viable yeast cells. (B) Transcriptional profiles of caspofungin-tolerant cells and micafungin-tolerant cells were highly similar to each other but distinct from either stationary or logarithmic controls.

In addition to performing RNAseq on samples enriched for tolerant cells, we also performed “bulk” RNAseq on unsorted echinocandin-treated cells as follows. 0.06 µg/ml of micafungin was added to 10^7^ *C. glabrata* cells/ml, which were then cultured for 24h, and their RNA was isolated and analyzed by RNAseq. Cells grown for 24h in the absence of micafungin (i.e., stationary phase cells) were used as the control.

### Tolerant cell-enriched RNAseq identified a unique transcriptional signature of echinocandin-tolerant cells

The sorted PI-negative cells were briefly treated with zymolyase to digest the fungal cell walls, followed by analysis using the single cell RNAseq pipeline at the Columbia University Genome Center. As is evident from the resulting UMAP schematic, the transcriptional profiles of caspofungin-tolerant cells and micafungin-tolerant cells were highly similar to each other but distinct from either stationary or logarithmic controls (Figure 1B). Therefore, for all following analyses, transcriptional data from caspofungin- and micafungin-tolerant cells were grouped together. Next, raw tolerant cell-enriched RNAseq data were normalized using the sctransform package and Seurat was used to generate a dataset of individual genes’ expression changes in echinocandin-tolerant cells relative to either stationary or logarithmic control cells (Dataset 1). We defined differentially expressed genes (DEGs) as showing at least one log2 unit (two-fold) expression difference in echinocandin-tolerant cells relative to no drug controls. DEGs were also defined in the “bulk” RNAseq dataset using the same criteria.

A comparison of “bulk” and “enriched” RNAseq datasets revealed a number of similarities and differences (Figure 2A, Figure S1A, B). Functional enrichment analysis (FUNCAT) showed that downregulated genes in all three datasets (“bulk” relative to stationary phase controls, “enriched” relative to stationary controls, and “enriched” relative to log phase controls) belonged to multiple functional categories involved in energy production and metabolism, particularly those related to mitochondrial functions (e.g., electron transport, respiration, and oxidative stress response) (Figure S1A-C). In contrast, there was no overlap between the FUNCAT categories of upregulated genes in “bulk” vs “enriched” RNAseq datasets (Figure S1A vs S1B, C). The “bulk” RNAseq dataset contained only a few upregulated FUNCAT categories, all of which related to cell wall metabolism (Figure S1A), consistent with the role of echinocandins in inhibiting cell wall biogenesis. In contrast, both “enriched” RNAseq datasets contained a larger number of upregulated FUNCAT gene categories, including ATP binding, DNA binding, DNA repair, DNA topology, and modification of chromosome structure (Figures S1B, C). Interestingly, although several individual cell wall maintenance genes were upregulated in the “enriched” datasets (see below), the cell wall maintenance FUNCAT categories were not identified in these datasets, suggesting that enriching for surviving cells does not necessarily enrich for cells with transcriptionally upregulated cell wall integrity pathways.

**Figure 2.**
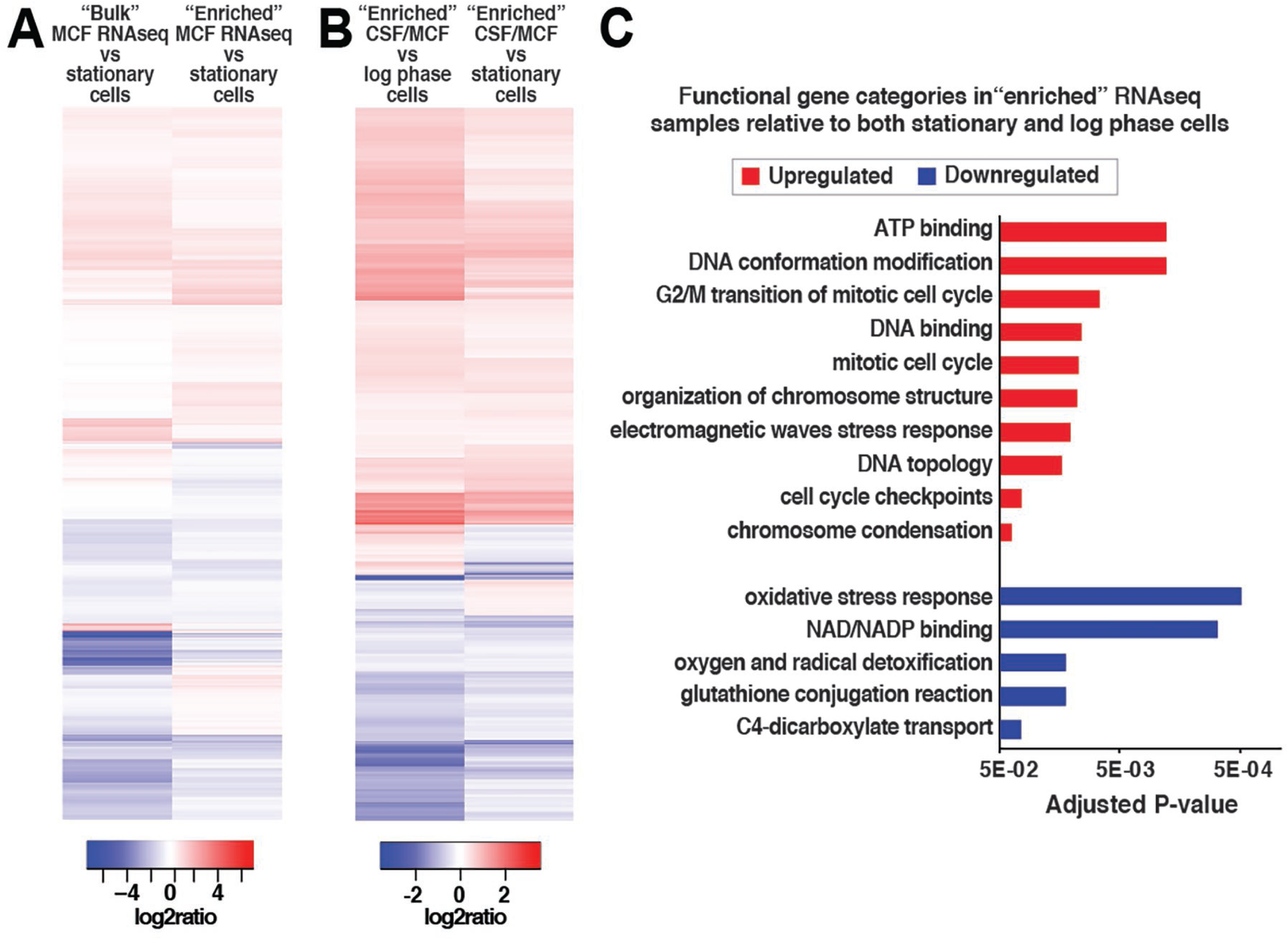
Echinocandin-tolerant *C. glabrata* cells upregulate genes involved in DNA topology and chromosome structure and downregulate genes involved in oxidative stress responses. (A) Heatmap showing the log2 ratios of “bulk” vs. tolerant cell-enriched (“enriched”) RNAseq samples. (B) Heatmap showing the log2 ratios of “enriched” RNAseq samples relative to “no drug” log phase or stationary phase controls. (C) Genes upregulated relative to both “no drug” controls were enriched for functional categories of ATP binding, DNA conformation modification and topology, and chromosome organization, whereas downregulated genes were enriched for functional categories of oxidative stress response, NAD/NADH binding, and oxygen and radical detoxification.

For the following analyses we focused on the RNAseq data obtained from samples enriched for echinocandin-tolerant cells. Comparisons of echinocandin-tolerant cells to either log phase or stationary phase controls resulted in similar but not identical expression patterns and FUNCAT categories (Figure 2B, Figures S1B, C), as is also evident from U-map analysis (Figure 1B). We were particularly interested in the genes that were upregulated or downregulated relative to both no drug controls (Dataset S1). The upregulated gene set was enriched for functional categories of ATP binding, DNA conformation modification and topology, and chromosome organization (Figure 2C). Consistent with echinocandins targeting β-1,3-glucan synthase (GS), an enzyme essential for maintaining cell wall structure, several genes involved in cell wall biosynthesis were upregulated in echinocandin-tolerant cells, including *FKS1* and *FKS2*, which encode GS, as well as *KRE5*, which encodes a protein required for β-1,6 glucan biosynthesis. Interestingly, several genes encoding multidrug ABC transporters (*CDR1* and *YOR1*) were also among the upregulated group, even though multidrug transporters are not thought to mediate resistance to echinocandins (18, 19). Genes downregulated relative to both no drug controls were enriched for functional categories of oxidative stress response, NAD/NADH binding, and oxygen and radical detoxification (Figure 2C). Of note, this is distinct from the stereotypical yeast environmental stress response, where these functional gene categories are typically upregulated (20, 21), underscoring that the transcriptional status of echinocandin-tolerant cells is unique and distinct from other types of stressed cells.

### Mitochondrial inhibitors but not topoisomerase inhibitors increase echinocandin tolerance in *C. glabrata*

Based on the functional categories of upregulated and downregulated genes, we selected different types of chemical inhibitors to modulate the most significantly up- or downregulated pathways and ask whether this modulation altered *C. glabrata* echinocandin tolerance. We note that, because caspofungin- and micafungin-tolerant cells had similar transcriptional profiles (Figure 1B), the experiments described below, especially the most laborious ones, were performed using only caspofungin. First, we treated *C. glabrata* with inhibitors of topoisomerase I (topotecan) or II (doxorubicin and etoposide) together with caspofungin for 24 hours. Etoposide and doxorubicin were used at concentrations corresponding to one third and one quarter of their minimal inhibitory concentration (MIC) in *C. glabrata*, respectively. Topotecan has been described to have no antifungal effect and not to affect viability, so we used 100 µM as did previous studies (22, 23) (Table S2). We found that addition of these topoisomerase inhibitors to cells cultured in the presence of caspofungin did not alter the *C. glabrata* tolerance phenotype (Figure 3A).

**Figure 3.**
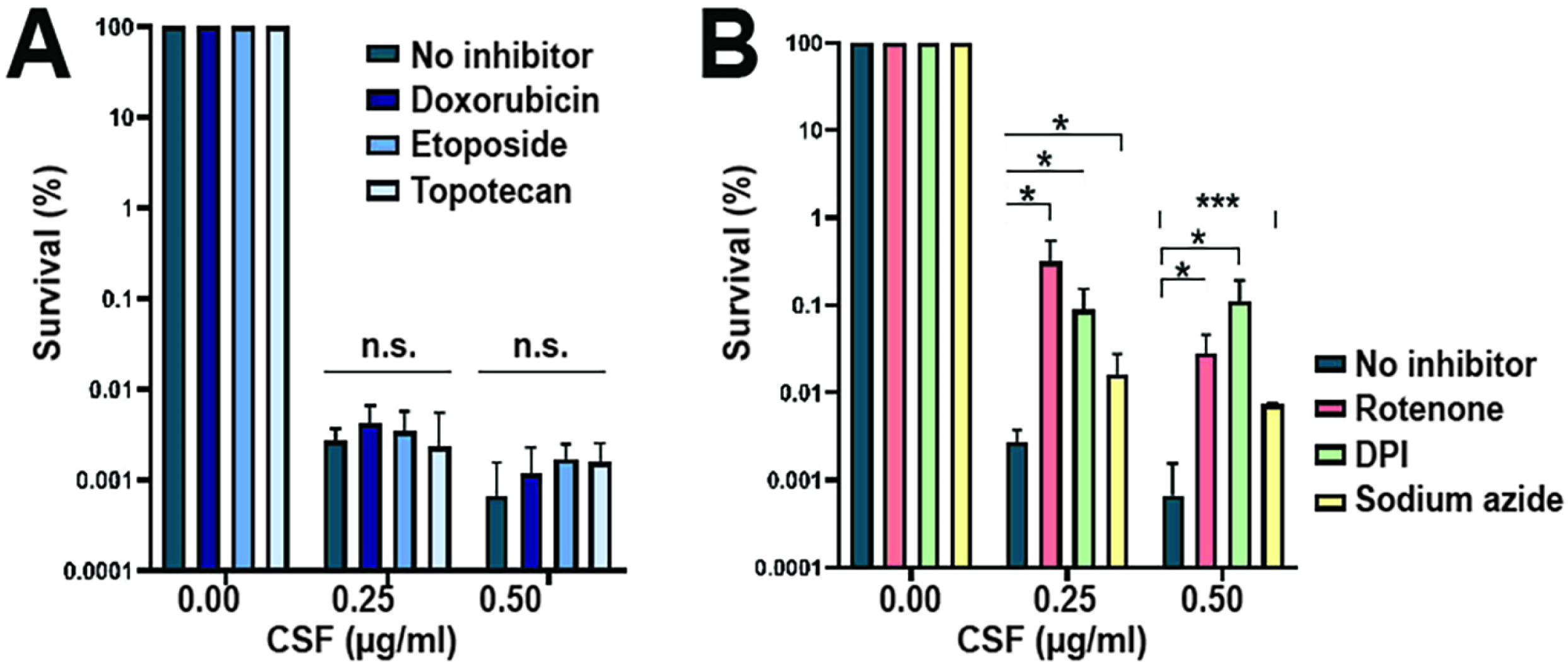
Mitochondrial inhibitors but not topoisomerase inhibitors increase echinocandin tolerance in *C. glabrata*. Chemical inhibitors were used to modulate the most significantly up- or downregulated pathways and examine the resulting effects on echinocandin tolerance. (A) Topoisomerase inhibitors did not alter echinocandin tolerance in *C. glabrata*, whereas (B) mitochondrial inhibitors increased the fraction of caspofungin-surviving cells. Statistical significance was calculated using an unpaired t-test and is indicated with asterisks (* p-value ≤0.05, ** p-value ≤0.01, *** p-value ≤0.001) or with n.s. (not significant).

Next, we asked whether inhibitors of mitochondrial enzymes that alter reactive oxygen species (ROS) production would alter tolerance to caspofungin. We used inhibitors of mitochondrial complex I (rotenone and diphenyleneiodonium chloride (DPI)) and complex IV (sodium azide) at concentrations corresponding to one third of their MIC in *C. glabrata* for DPI and sodium azide. The rotenone concentration used (0.3 mM) was determined by the highest concentration at which it was soluble in YPD medium (Table S2). We observed that these inhibitors increased the fraction of cells surviving after 24 hours of treatment by 10 to 100-fold (Figure 3B). Both complex I inhibitors (rotenone and DPI) improved survival more than complex IV inhibitor sodium azide.

### Caspofungin treatment leads to an increase in reactive oxygen species production but a downregulation of the oxidative stress response

Increased ROS have been observed as a consequence of drug treatment both in fungi and bacteria and have been proposed as a key mediator of drug-induced killing in some of these systems (24–26). This, together with the observation that mitochondrial inhibitors improve echinocandin tolerance, led us to examine ROS levels and ROS responses in caspofungin-treated cells more closely. A typical method to detect ROS is by using ROS-sensitive dyes followed by flow cytometry analysis. We used two different dyes, 2′,7′-dichlorodihydrofluorescein diacetate (CFDA) and dihydroethidium (DHE), which are thought to detect specific ROS: hydrogen peroxide and superoxide, respectively (27). However, the exact nature of the species detected by these dyes, especially CFDA, is still obscure (28). In these experiments, *C. glabrata* cells were treated with the indicated agents for 2, 6, or 24 hours, followed by staining with PI or Sytox Green to gate out dead cells and CFDA or DHE to detect ROS (PI was used with CFDA and Sytox Green was used with DHE due to color compatibility). We used treatment with 50 mM H_2_O_2_ as a positive control and an echinocandin-resistant *fks1* mutant as a negative control. As expected, we detected a strong increase in both CFDA and DHE staining induced by H_2_O_2_ treatment, albeit with different dynamics: CFDA staining peaked at 2 hours of exposure and then declined over 24 hours, whereas DHE staining gradually increased over 24 hours (Figure 4A). Importantly, we observed that 0.25 µg/ml caspofungin also induced a strong increase in both CFDA and DHE staining, with staining intensity of both dyes increasing over the 24-hour period. This was entirely dependent on the presence of wild type GS enzyme, as no increase was observed in the *fks1* mutant (Figure 4A). Thus, we concluded that caspofungin-mediated inhibition of GS activity resulted in an increase in cellular ROS levels.

**Figure 4.**
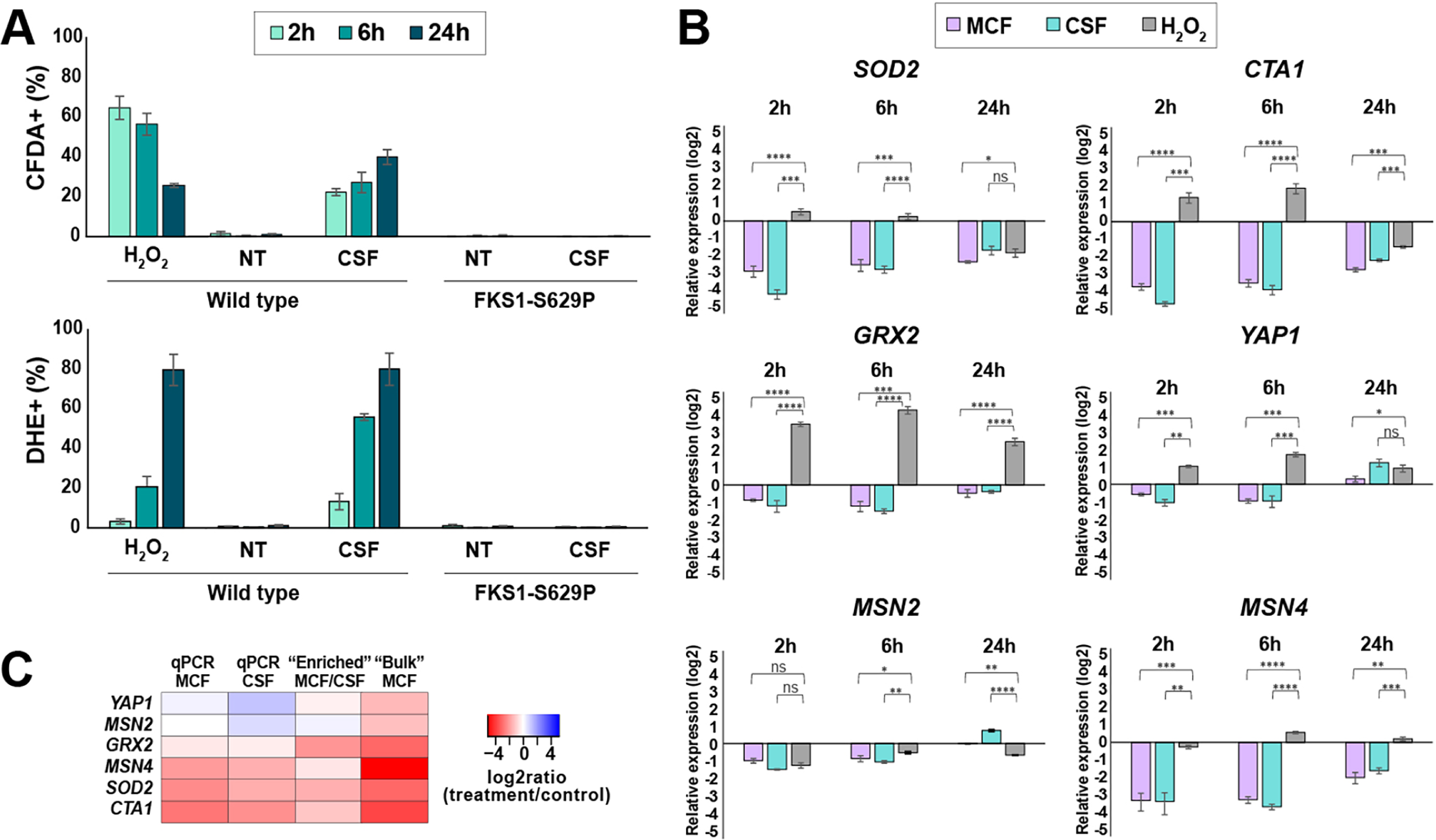
Caspofungin treatment leads to increased reactive oxygen species (ROS) levels but decreased expression of genes involved in oxidative stress responses. (A) Two different ROS-sensitive dyes, CFDA or DHE, were used to detect hydrogen peroxide or superoxide species, respectively. Dead cells were gated out using PI or Sytox Green (SG). 50 mM H_2_O_2_ as a positive control. Caspofungin induced a strong increase in both CFDA and DHE staining, and this increase was abolished in an *fks1* mutant, which lacks wild type glucan synthase (echinocandin target enzyme). NT indicates caspofungin non-treated cells. (B) Oxidative stress response (OSR) genes as well as transcription factors involved in the regulation of the OSR pathway were downregulated by caspofungin and micafungin treatment but were either induced or unaltered by the H_2_O_2_ control. (C) Heatmap comparing log2 ratios of treatment/control obtained using different molecular approaches for the six selected genes involved in oxidative stress responses. The statistical significance was calculated using an unpaired t-test and is indicated with asterisks (* p-value ≤0.05, ** p-value ≤0.01, *** p-value ≤0.001, **** p-value ≤0.0001) or with ns (not significant).

The observed increase in ROS but a decrease in the expression of genes involved in oxidative stress responses (Figure 2C, Figure S1) by RNA-seq was surprising, so we probed it further by examining the transcriptional levels of several individual genes during caspofungin or micafungin treatment by quantitative reverse transcriptase PCR (qRT-PCR). These experiments were performed by harvesting “bulk” RNA from echinocandin-treated or untreated cultured, followed by qPCR analysis. We chose genes with well-established roles in the oxidative stress response (OSR), such as mitochondrial manganese superoxide dismutase (*SOD2*, *CAGL0E04356g*), a catalase A (*CTA1*, *CAGL0K10868g*) and a glutathione oxidoreductase (*GRX2*, *CAGL0K05813g*), as well as three transcription factors that regulate the OSR pathway (*YAP1*, *CAGL0H04631g*; *MSN2*, *CAGL0F05995g*; and *MSN4*, *CAGL0M13189g*) (29). The relative expression of each gene was normalized to a control locus (*RDN5.8*) and calculated relative to that of untreated stationary phase cells. H_2_O_2_ treatment was used as a control. Consistent with the RNAseq results (Figure 2C), we observed that transcript levels of all these genes were downregulated by caspofungin and micafungin treatment, with *SOD2*, *CTA1*, and *MSN4* showing the greatest reduction in RNA abundance (up to 4 log2 units, or 16-fold, Figure 4B). This transcriptional response was highly distinct from that induced by H_2_O_2_, which predominantly resulted either in an increase or no change in these genes’ transcript abundance (Figure 4B). The log2 ratios (treatment/no treatment) obtained for these genes by qPCR showed good agreement with “bulk” and “enriched” RNAseq results (Figure 4C).

### ROS reduction per se does not increase echinocandin tolerance in *C. glabrata*

As mentioned above, ROS are generally viewed as cell damaging agents and have been shown to mediate the killing effects of certain antimicrobial drugs. To investigate whether this is the case for echinocandins in *C. glabrata*, we turned our focus again to the mitochondrial inhibitors. First, we examined their effects on caspofungin-induced killing of *C. glabrata* over a wide range of caspofungin concentrations (Figure 5A). We believe such killing assay is a better indicator of drug tolerance than the standard minimum inhibitory concentration (MIC) determination, which measures growth but not survival in the presence of drug (8). The caspofungin MIC-value obtained for ATCC2001 was 0.12 µg/ml, and it was not affected by any of the mitochondrial inhibitors (Table S2). Nevertheless, survival after 24 hours in caspofungin was rescued by rotenone, DPI and sodium azide at several different caspofungin concentrations (Figure 5A). Next, we examined how ROS production was affected by these mitochondrial inhibitors and found that they reduced ROS levels in caspofungin-treated cells (Figure 5B). This was especially evident when ROS was detected by CFDA staining but less so when DHE staining was used, suggesting that these compounds reduce the abundance of some ROS species more than others.

**Figure 5.**
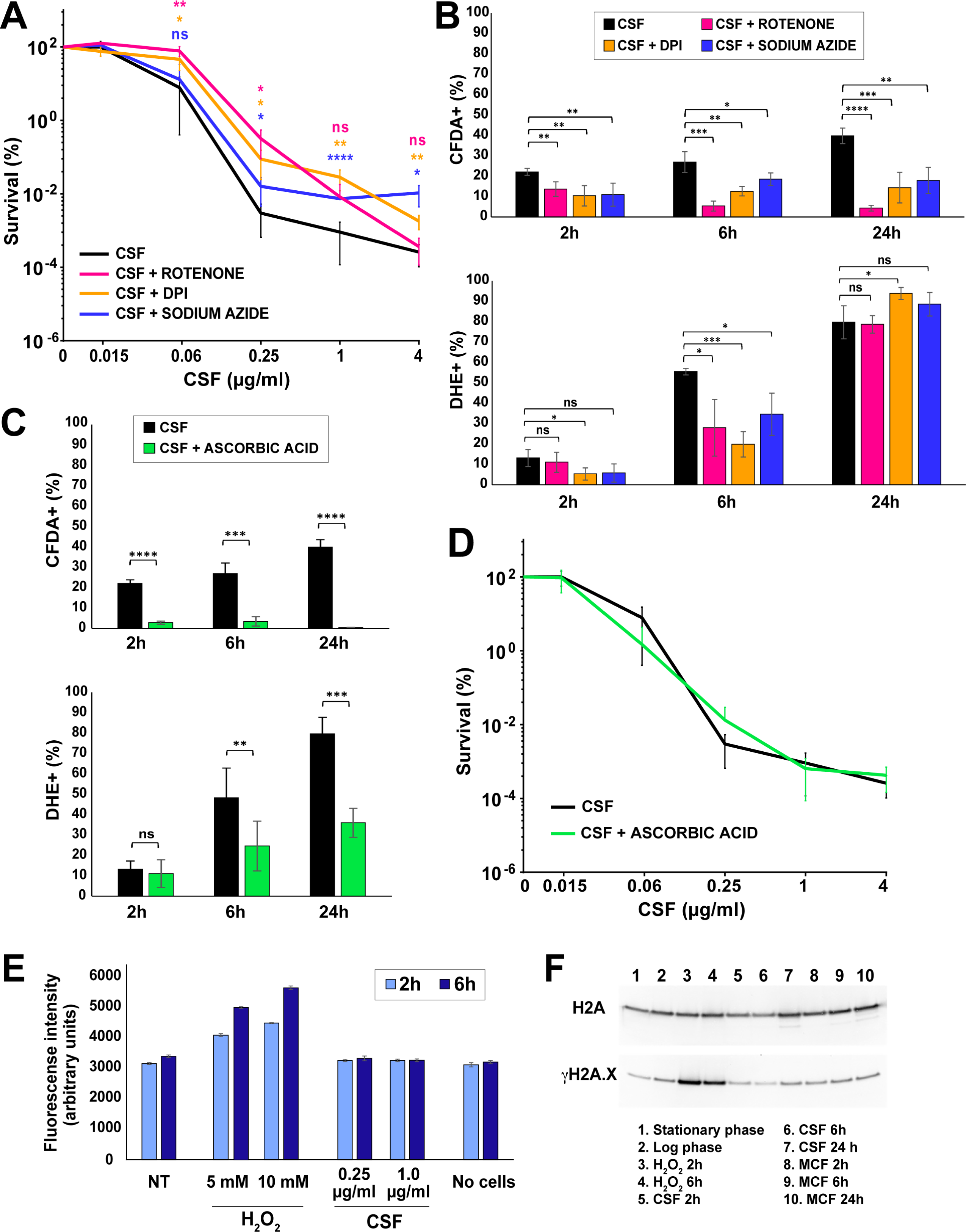
Mitochondrial inhibitors but not an ROS scavenger improve survival of *C. glabrata* cells in caspofungin. (A) Mitochondrial inhibitors rotenone, DPI and sodium azide increased the fraction of caspofungin-surviving cells over a wide range of caspofungin concentrations. (B) The mitochondrial inhibitors also reduced ROS production as detected by CFDA and DHE. (C) Ascorbic acid, an ROS scavenger, strongly decreased ROS levels induced by caspofungin. (D) Ascorbic acid did not alter cell survival in the presence of caspofungin. (E) Despite the elevation of ROS, no increase in lipid peroxidation, as measured by the DPPP fluorometric assay, was detected in the presence of caspofungin. (F) No increase in DNA damage as measured by γH2A.X abundance was detected in the presence of caspofungin or micafungin. The statistical significance was calculated using an unpaired t-test and is indicated with asterisks (* p-value ≤0.05, ** p-value ≤0.01, *** p-value ≤0.001, **** p-value ≤0.0001) or with ns (not significant).

The result above was consistent with the hypothesis that elevated ROS may contribute to echinocandin-induced cell death in *C. glabrata*. To test this hypothesis directly, we used ROS scavengers. We tested several compounds with ROS scavenging activities (thiourea, dimethylthiourea, sodium dithionite, catalase, α-tocopherol and ascorbic acid), but found that only ascorbic acid significantly reduced the levels of ROS in caspofungin-treated cells (Figure 5C). Interestingly, when we measured cell survival of caspofungin treatment in the presence of ascorbic acid, we found no difference between cells cultured in the absence or presence of this ROS scavenger (Figure 5D). This result indicated that it was not ROS per se that was responsible for cell death upon caspofungin treatment.

ROS can damage different cellular biomolecules, which is thought to contribute to their involvement in cell death in other contexts (30). Thus, we examined whether elevated ROS levels in caspofungin-treated cells were associated with detectable damage of two types of biomolecules: lipids and DNA. To measure lipid peroxidation, a fluorometric assay was performed using diphenyl-1-pyrenylphosphine (DPPP), a compound that reacts with lipid hydroperoxides to produce fluorescent DPPP oxide. To measure DNA damage, we examined the abundance of histone H2A phosphorylated at serine 129 (γH2A.X), which is a universal marker of double-strand breaks (31). In both cases, H_2_O_2_ was used as a positive control. Interestingly, whereas H_2_O_2_-treated cells showed robust induction of lipid peroxidation and γH2A.X levels, echinocandin-treated cells did not show evidence of either lipid peroxidation or DNA damage relative to non-treated controls (Figure 5E, F). This result was consistent with the conclusion that despite an increased abundance of ROS in echinocandin-treated cells, these ROS did not significantly contribute to cellular damage or lethality.

### Deletion mutants lacking components of the mitochondrial respiratory chain show a caspofungin hypo-tolerant phenotype

The observations that mitochondrial inhibitors altered echinocandin tolerance but that ROS per se were not involved made us examine the role of mitochondria in echinocandin tolerance more closely. To this end, we generated a number of different mitochondria-deficient strains (Table S3). First, we created deletions of five genes encoding components of the mitochondrial respiratory chain: *NDI1* (*CAGL0B02431g*), whose *Saccharomyces cerevisiae* ortholog has NADH dehydrogenase activity and is a component of the fungal equivalent of mitochondrial complex I; *COX4* (*CAGL0L06160g*), whose ortholog has the cytochrome-c oxidase activity of mitochondrial complex IV, and three components of the mitochondrial complex V ATP synthase – the F1 alpha subunit of the F1F0-ATPase complex encoded by *ATP1* (*CAGL0M09581g*), the F1 beta subunit encoded by *ATP2* (*CAGL0H00506g*), and the assembly factor for the F0 sector of mitochondrial F1F0 ATP synthase encoded by *ATP10* (*CAGL0C02651g*). All these deletion mutants, with the exception of *ndi1Δ*, showed a petite phenotype characterized by slow growth and inability to use a non-fermentable carbon source (Figure S2). We also isolated several respiration-deficient (a.k.a. petite) mutants that spontaneously but rarely (approximately one in 3000-5000 colonies) emerged after 24 hours of 0.25 µg/ml caspofungin treatment. Finally, we generated several petite mutants by treatment with ethidium bromide (32). All isolated petite mutants were respiration-deficient and slow growing; however, all of them stained to various degrees by MitoTracker Green, indicating that they were not fully lacking mitochondria (Figure S3A). As expected, most mitochondrial mutants, including several gene deletions (*atp1Δ, atp2Δ*, and *atp10Δ*) and all petite mutants, showed elevated MICs to fluconazole (Table S3), consistent with previous reports that mitochondrial dysfunction in *C. glabrata* induces *PDR1* expression and promotes azole resistance (33, 34).

All mitochondria-deficient mutants generated above were examined in the caspofungin killing assay. Interestingly, and contrary to the expectation formed based on the phenotype of mitochondrial inhibitors, all deletion mutants (including *ndi1Δ*, which was not petite) showed a caspofungin hypo-tolerant phenotype at one or more concentrations (Figure 6A, B). The same hypo-tolerant phenotype was observed in all examined petite mutants (Figure 6C and Figure S3B). Because this hypo-tolerance was different from the phenotype caused by inhibitors of the mitochondrial respiratory chain, we wondered whether it may be due not to a lack of respiratory genes per se but to general mitochondrial defects. To test this, we also deleted three genes encoding mitochondrial proteins not directly involved in the respiratory chain: *PET9* (*CAGL0F04213g*), encoding major ADP/ATP carrier of the mitochondrial inner membrane; *TIM18* (*CAGL0A03784g*), encoding a translocase of the inner mitochondrial membrane; and *YME1* (*CAGL0K05093g*), encoding a catalytic subunit of i-AAA protease complex responsible for degradation of misfolded mitochondrial gene products. We found that each of these mutants was likewise hypo-tolerant to caspofungin, indicating that this hypo-tolerance may be a general feature of *C. glabrata* strains with mitochondrial defects (Figure S3C).

**Figure 6.**
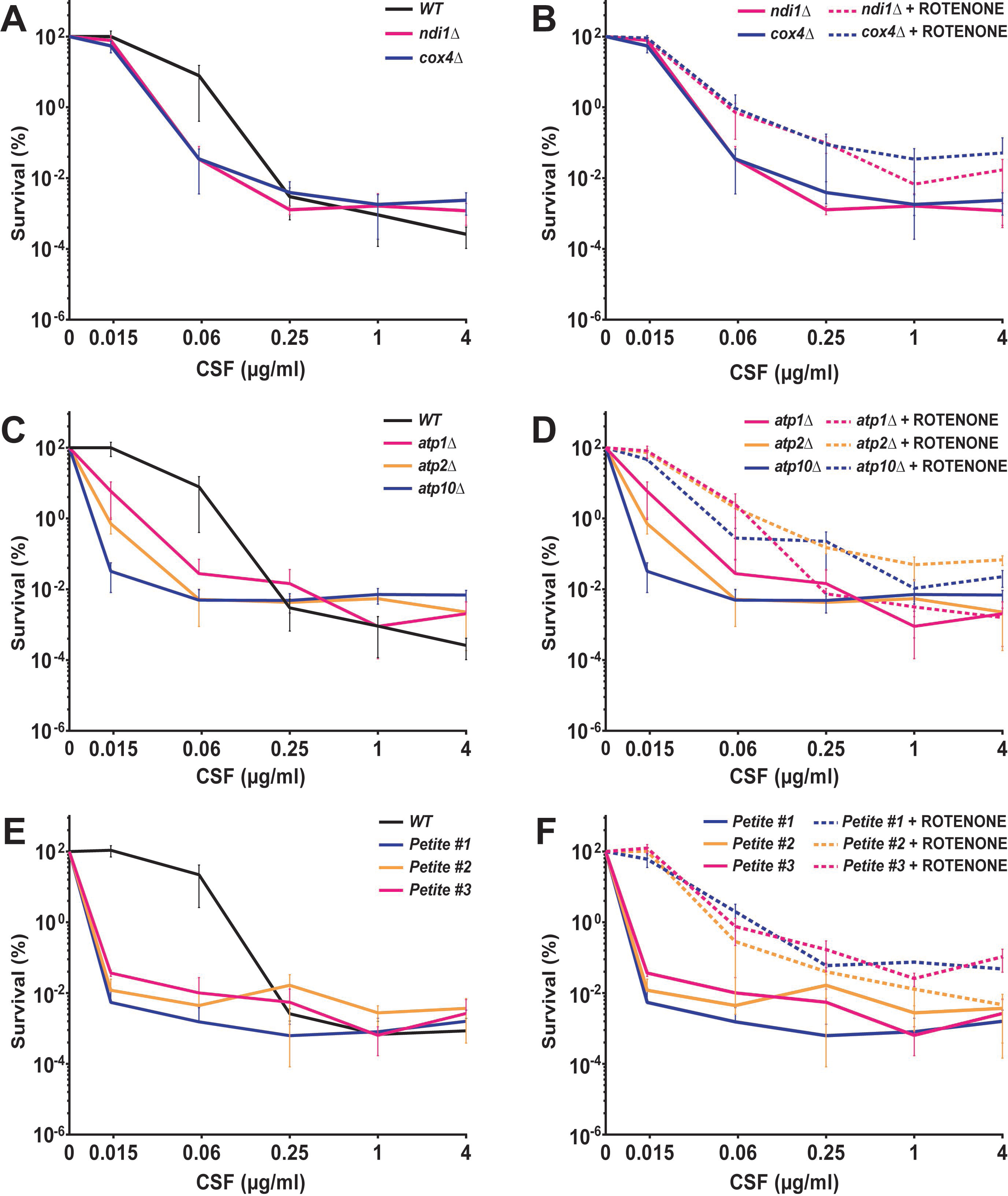
Mutants lacking various mitochondrial respiratory chain components and mitochondrial-defective petite mutants showed a caspofungin hyper-susceptible phenotype, which was rescued by rotenone. (A) Deletion mutant of mitochondrial respiratory chain component *NDI1* (NADH dehydrogenase, the yeast equivalent of complex I) and *COX4* (cytochrome-c oxidase of complex IV) showed a mild hyper-susceptibility to caspofungin that was rescued by rotenone (B). (C) Deletion mutants of three components of the mitochondrial complex V ATP synthase (*ATP1*, *ATP2* and *ATP10*) also showed a hyper-susceptible phenotype to a wide range of caspofungin concentrations below the MIC-value that was rescued by rotenone (D). (E) Respiration-deficient petite mutants generated during caspofungin exposure showed the strongest hyper-susceptible phenotypes that were also rescued by rotenone (F).

Interestingly, the hypo-tolerance of mitochondrial mutants to caspofungin was rescued by rotenone in every tested case (Figure 6D, E, F), indicating that rotenone’s effect on echinocandin tolerance is exerted whether cells have functional mitochondria or not. In contrast, DPI did not improved survival in any examined mitochondrial mutants, indicating that its effect was mediated by mitochondria (Figure S4A, S4B). Furthermore, we examined the effect of sodium azide on the *cox4Δ* mutant because complex IV is the target of this mitochondrial inhibitor. No rescue of survival was observed, indicating that a functional complex IV is necessary to mediate the effect of sodium azide (Figure S4C).

To test whether the hypo-tolerance to caspofungin was due to a general cell wall integrity defect, we assayed the sensitivity of mitochondrial mutants to two different cell wall damaging agents: Congo red and calcofluor white. We found that none of the examined mutants were more sensitive to these agents than the parental wild type strains (Figure S2), indicating that the hypo-tolerance effect was specific to echinocandin-mediated cell killing.

Because the effect of mitochondrial inhibitors (which increased tolerance) was opposite of that of mitochondrial mutants (which reduced tolerance), we asked whether one important difference between the two was the timing of the onset of mitochondrial dysfunction. In the experiments using mitochondrial inhibitors, the inhibitors had been added to cells with intact mitochondria at the same time as the echinocandins. However, in the mutants, the mitochondria had been compromised by mutation prior to addition of echinocandins. Therefore, we asked whether inhibition of mitochondria prior to echinocandin exposure would elicit different effects relative to mitochondrial inhibition induced at the same time as echinocandin exposure. We cultured *C. glabrata* for 18 hours in the presence of rotenone or DPI, then we collected the cells and cultured in fresh YPD media with the mitochondrial inhibitor and caspofungin for another 24 hours, as in the standard tolerance assay (thus, rotenone or DPI were present in the culture both before and after caspofungin addition). Interestingly, pre-exposure of *C. glabrata* to DPI abolished the rescuing effect of DPI on caspofungin tolerance at all tested concentrations, and this was also true for rotenone at one sub-MIC caspofungin concentration (0.06 µg/ml) (Figure S5). These results suggest that chronic mitochondrial inhibition or dysfunction reduces *C. glabrata* echinocandin tolerance, whereas acute mitochondrial inhibition may enhance tolerance.

### Glucan synthase enzyme from mitochondrial mutants exhibits normal echinocandin sensitivity *in vitro*

Echinocandins kill fungal cells by targeting the enzyme 1,3-β-D-glucan synthase (GS) and inhibiting fungal cell wall formation. Thus, the increased echinocandin susceptibility of mitochondrial mutants could potentially be due to changes in GS, which would make the enzyme less susceptible to drug action, as we had demonstrated with echinocandin resistance in *Aspergillus fumigatus* (35), or to changes in pathways acting downstream of GS inhibition. To distinguish between these possibilities, we partially purified GS complexes using our established method consisting of membrane protein extraction and partial GS purification by product entrapment (36) from two different mitochondrial mutants – the *ndi1Δ* strain and one petite mutant – and a wild type control strain. The activity of the isolated GS complexes and their inhibition by micafungin were assayed *in vitro* using a β-glucan polymerization assay (36). We observed that GS isolated from mitochondrial mutants had very similar inhibition kinetics to GS isolated from the wild type strain (Figure 7), indicating that the effect of mitochondria on echinocandin sensitivity likely occurs downstream of the GS inhibition step.

**Figure 7.**
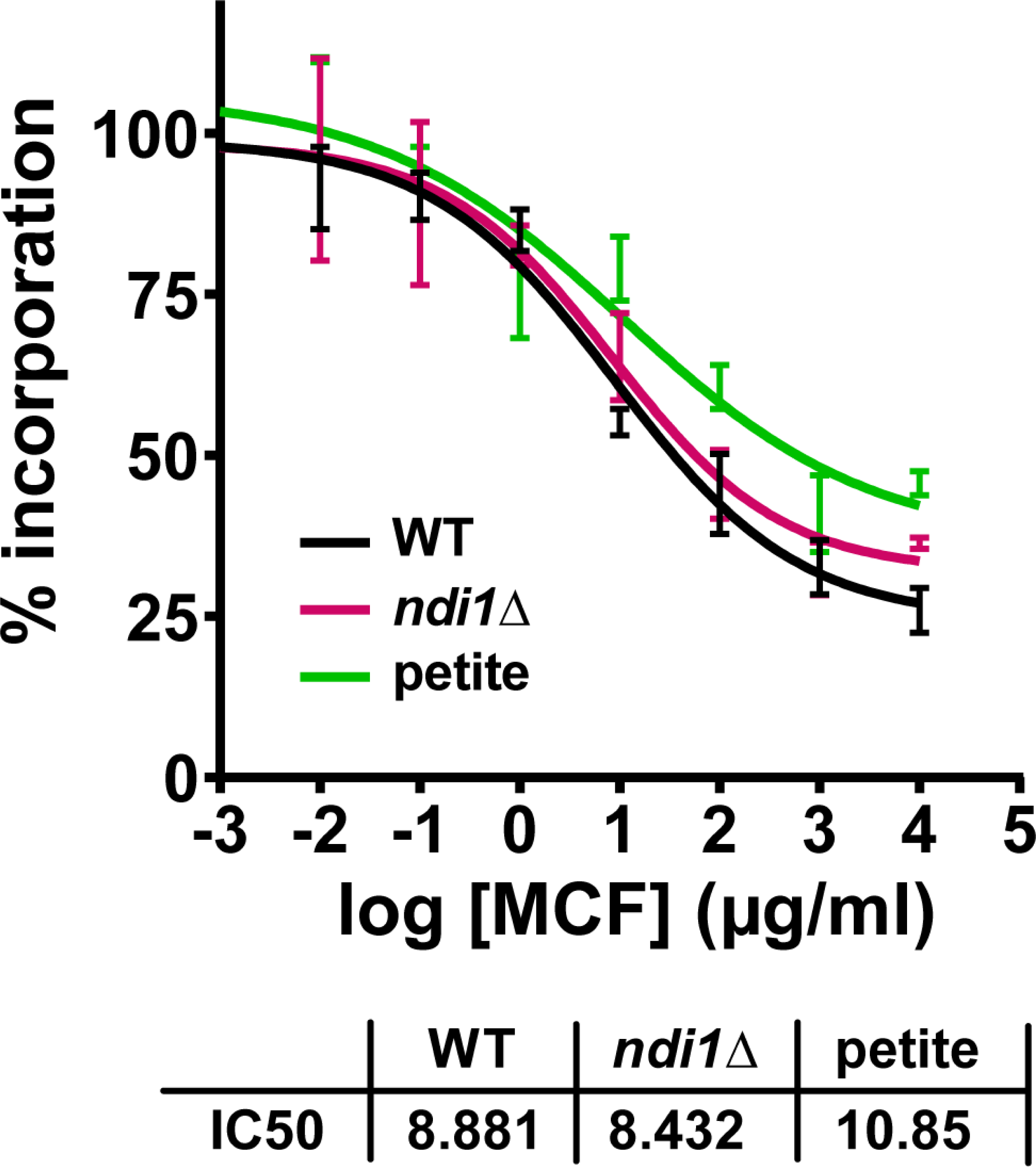
1,3-β-D-glucan synthase (GS) isolated from mitochondrial mutants shows unaltered micafungin inhibition kinetics *in vitro*. Echinocandin inhibition profiles and biochemical sensitivity (IC50) for product-entrapped GS enzyme complexes was assessed by the incorporation of [^3^H]glucose into radiolabeled product. Micafungin titration curves and EC50 values are shown for two different mitochondrial mutants – the *ndi1Δ* strain and one petite mutant – and the wild type control strain.

## DISCUSSION

Our study presents the first examination of the transcriptional landscape of the echinocandin-tolerant cell subset in the human pathogen *C. glabrata*, which is presumed to serve as a precursor for stable drug resistance (8, 11). We show that the transcriptional state of cells that have survived 24 hours of above-MIC echinocandin exposure is distinct from the stereotypical transcriptional response to environmental stresses. In particular, echinocandin-tolerant cells upregulated pathways involved in chromosome structure and DNA topology and downregulated pathways involved in oxidative stress response. This downregulation occurred even though *C. glabrata* cells had significantly elevated ROS levels upon echinocandin exposure. Interestingly, the ROS did not appear to contribute to echinocandin-induced cell death, as reducing ROS using ascorbic acid did not alter cell killing. Finally, we identified a multifactorial role of mitochondria in *C. glabrata* echinocandin tolerance: whereas using inhibitors of respiratory chain components increased echinocandin tolerance, deletion of respiratory chain components made *C. glabrata* cells hypo-tolerant to echinocandins. Together, these results provide new insights into the *C. glabrata* response to echinocandins and reveal the involvement of mitochondria in echinocandin tolerance. A model describing the proposed role of mitochondria in echinocandin-induced cell death is shown in Figure 8.

**Figure 8.**
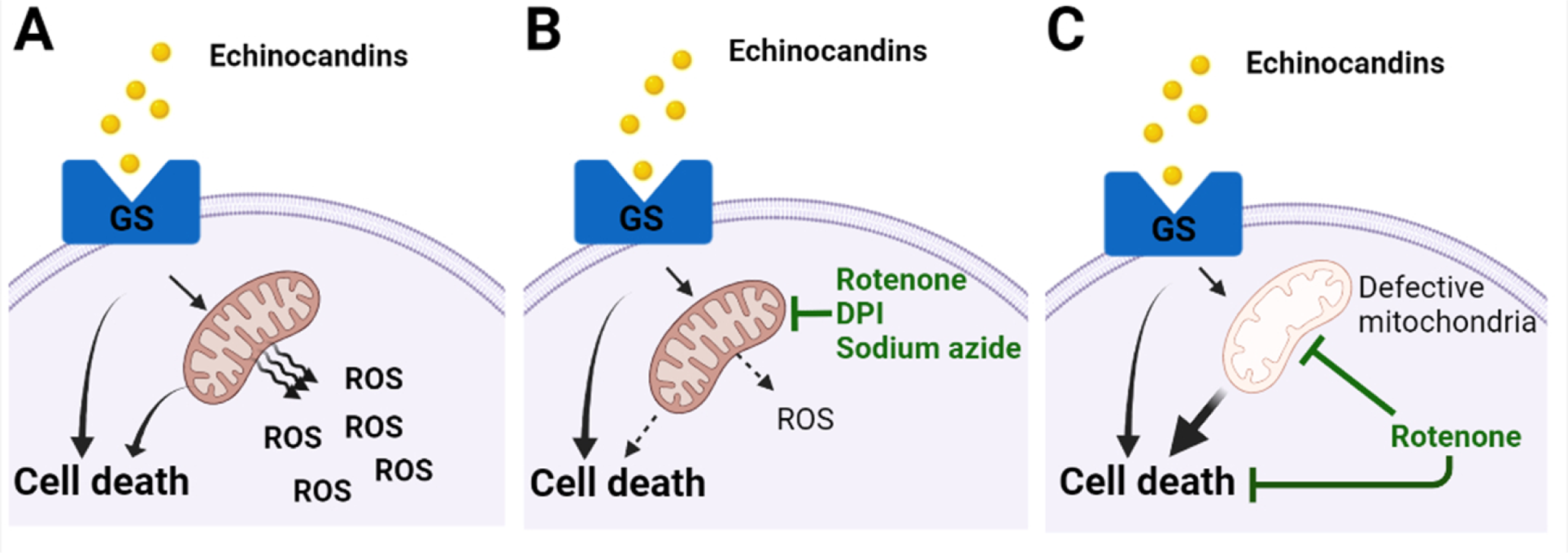
Graphical representation of the proposed model for the role of mitochondria in echinocandin tolerance. (A) Echinocandin exposure in *Candida glabrata* cells inhibits GS activity, leading to cell death via mitochondria-dependent as well as mitochondria-independent mechanisms. The mitochondrial involvement also results in increased ROS production. (B) When echinocandin exposure is combined with mitochondrial inhibitors, mitochondria-dependent mechanisms of cell death are reduced, along with ROS production. (C) Chronic mitochondrial dysfunction leads to hypo-tolerance to echinocandins characterized by decreased survival at below-MIC concentrations. In this case, rotenone still partially rescues survival, likely due to non-mitochondrial mechanisms. Created with BioRender.com

Several gene categories upregulated in echinocandin-tolerant cells were involved in DNA conformation modification and chromosome organization. However, using topoisomerase inhibitors did not alter the killing dynamics of echinocandin drugs, suggesting that the pathways may not contribute to survival but instead serve other roles in tolerant cells. *C. glabrata* is known for its genome plasticity: its clinical isolates show remarkable genetic diversity both in terms of nucleotide sequence and chromosome structure (37–39). However, it is not known under what circumstances these genomic changes are acquired, as cells cultured in rich medium in the lab appear to be karyotypically stable (37). Thus, cells dealing with long-term echinocandin-induced stress may activate mechanisms enabling chromosomal variation to enhance adaptability. In this context, it is also interesting that we did not detect an increase in abundance of the “DNA damage histone” γH2A.X during echinocandin exposure, suggesting that DNA double-strand breaks (which induce γH2A.X formation) either may not be the main lesions triggering chromosomal alterations, or that they occur at lower levels than can be detected by Western blotting. Nevertheless, the transcriptional upregulation of DNA topology and chromosomal conformation genes suggests that these cells may be in a specialized activated transcriptional state that resists drug stress and has an elevated genetic potential to evolve into a genotypically stable drug-resistant cell. Further studies addressing the development of resistant mutations in echinocandin tolerant cells pre-exposed to this drug class will shed light on this issue.

Most transcriptional studies of fungal species exposed to drugs have been performed using short-term treatments and traditional bulk RNA-seq approaches with no enrichment for surviving cells (15, 16, 40). Because echinocandins rapidly induce cell death (17), such bulk analysis performed after a few hours of treatment overwhelmingly includes dead and dying cells. In contrast, our RNA-seq approach was designed to enrich for the rare *C. glabrata* cells surviving 24 hours of above-MIC drug exposure. Thus, our approach identified a distinct transcriptional profile of echinocandin-tolerant cells compared to other studies. Because the echinocandins target cell wall assembly, previous analyses of echinocandin-induced transcriptional changes mainly focused on cell wall-related processes. For instance, Xiong et al. described two *C. albicans* transcription factors regulating cell wall maintenance and homeostasis (Efg1 and Cas5) involved in the transcriptional response to two-hour caspofungin treatment (15). Likewise, in *Aspergillus fumigatus* it has been reported that caspofungin exposure leads to communication between the cell wall-integrity and the high osmolarity-glycerol pathways through MAPK signaling (41). Other studies have also identified transcription factors that play a role in the regulation of the fungal cell wall as well as in echinocandin tolerance (40). Whereas our study also identified several upregulated cell wall maintenance genes in echinocandin-tolerant cells, including *FKS1* and *FKS2*, the cell wall integrity pathway as a whole was not present among the upregulated FUNCAT categories, suggesting that during long-term echinocandin exposure *C. glabrata* cells may switch to other strategies to survive.

In most examined human-pathogenic yeasts and molds, amphotericin B and azoles trigger intracellular ROS formation, which contributes to the antifungal action of these drugs (24, 25, 42, 43). Echinocandins are also associated with an increase in ROS production (25, 43, 44) and, at least in *C. albicans*, ROS scavenger thiourea caused a decrease in micafungin-induced killing (43). Also in *C. albicans*, genes involved in oxidative stress responses are induced in response to caspofungin (44). Our results (both RNA-seq and qRT-PCR of individual oxidative stress responder and regulator genes) show that although echinocandins also induce robust ROS formation in *C. glabrata*, this fungus does not respond to echinocandins by upregulating oxidative stress response pathways. Indeed, the high ROS levels might be a consequence of the downregulation of expression of the ROS detoxifying enzymes. Furthermore, we could not elicit a reduction of ROS using thiourea or several other scavengers, with the exception of ascorbic acid, which strongly reduced ROS levels in echinocandin-treated cells. Despite this decrease, however, ascorbic acid did not alter the dynamics of cell killing, indicating that in *C. glabrata* the ROS do not significantly contribute to cell death but rather serve another function. Consistent with this idea, the ROS increase was not associated with detectable lipid peroxidation or DNA damage. Although these results run contrary to the traditional view that ROS are exclusively damaging and lack other physiological functions, they are consistent with observations that *C. glabrata* is exceptionally resistant to oxidative stress (45) and in fact, it has recently been described that petite mutants of *C. glabrata* show a lack of oxidative stress susceptibility when incubated with hydrogen peroxide which is combined with a constitutive upregulation of environmental stress-response and heat-shock-protein-encoding genes (46). Relative to other species, *C. glabrata* exhibit a very restrained transcriptional response to phagocyte-induced ROS with insignificant upregulation of catalase (47). Interestingly, recent evidence suggests a critical role of ROS in the communication between the mitochondria and other cellular processes to maintain homeostasis and promote adaptation to stress (48). Furthermore, some studies have also proposed a role for oxidative stress adaptation in promoting virulence and drug resistance in *C. glabrata*, as several genes implicated in virulence, biofilm formation, and drug transport were also induced during the oxidative adaptation response (45). These studies, together with our results, suggest that in *C. glabrata* responding to echinocandins ROS function as a signaling intermediate with as yet unknown specific downstream targets and contribution to fitness.

The involvement of mitochondria in echinocandin tolerance in *C. glabrata* was suggested by the RNA-seq data and the observation that mitochondrial inhibitors reduced *C. glabrata* killing by echinocandins over a range of concentrations. In contrast, however, deletion mutants with defective respiratory chain functions, deletions of several other mitochondrial genes, as well as a series of respiration-deficient petite strains of unknown etiology, were all hypo-tolerant to echinocandins, showing that mitochondrial inhibition and genetically encoded mitochondrial deficiency have opposite effects on echinocandin tolerance. Reduced tolerance to echinocandins in some mitochondrial mutants was previously described in *C. albicans* and *S. cerevisiae*. For instance, several *C. albicans* and *S. cerevisiae* mitochondrial mutants are sensitive to multiple cell wall damaging agents, including calcofluor white, Congo red, and echinocandins, indicating a general cell wall defect (49, 50). This defect has been attributed to the role of mitochondria in phospholipid metabolism, whereby mitochondrial defects compromise phospholipid production, affecting cell wall biogenesis (49, 51). Similarly, *S. cerevisiae* mutants defective in producing cardiolipin, a predominantly mitochondrial lipid, have general cell wall defects (52). Interestingly, none of our *C. glabrata* mitochondrial mutants showed increased sensitivity to Congo red or calcofluor white, indicating that the mitochondrial effect on the cell wall in *C. glabrata* may be limited to β-glucans. Possibly relatedly, in *S. cerevisiae*, abrogation of the cardiolipin biosynthetic pathway results in reduced β-glucan levels and can be rescued by mutation of *KRE5*, which encodes a β-1,6-glucan synthase (52). A direct link between glucan synthesis and plasma membrane lipid composition has also been demonstrated in *Aspergillus fumigatus*, where two sphingolipids, dihydrosphingosine and phytosphingosine, rendered GS insensitive to echinocandins (35). Thus, although our RNA-seq analysis did not reveal a lipid-related transcriptional signature in echinocandin-tolerant cells, the studies described above and our results obtained with mitochondrial mutants support a model wherein mitochondrial status may influence the plasma membrane lipid composition, which in turn may influence echinocandin sensitivity.

Another potential link between mitochondria and echinocandin sensitivity has been suggested by reports indicating that in *C. albicans* and in *C. glabrata* echinocandins induce cell death by apoptosis (programmed cell death) and by necrosis (53, 54). It is well known that the role of mitochondria in programmed cell death is complex, and that mitochondria have both pro- and anti-apoptotic functions (55). Thus, it may not be surprising that inhibition of certain mitochondrial activities leads to different survival phenotypes than a total lack of certain mitochondrial proteins due to gene deletions. Interestingly, our results suggest that at least one of the mitochondrial inhibitors, rotenone, which elicited the strongest rescue of cell survival, may not be acting solely via the mitochondria. Rotenone is typically considered a mitochondrial complex I inhibitor. However, *C. glabrata*, like some other fungal species, has lost the canonical NADH:ubiquinone oxidoreductase (complex I) while retaining other mitochondrial oxidative phosphorylation components, including NADH dehydrogenases (encoded by *NDI1*, *NDE1*, and *NDE2*), which largely perform the same function as complex I. The observation that rotenone improved survival not just in wild type cells, but also in every examined mitochondrial mutant, suggests that this rotenone effect is achieved not just via mitochondrial modulation but via influencing other, as yet unclear secondary pathways. To identify these pathways, transcriptomic studies will likely be highly informative. In contrast, DPI (an inhibitor of NADH dehydrogenases) and sodium azide (an inhibitor of complex IV) both failed to improve survival in mitochondrial mutants, suggesting that their activity is indeed mediated by the mitochondria. Surprisingly, the transcriptome of *C. glabrata* echinocandin tolerant cells showed a downregulation of multiple genes involved in mitochondrial function, e.g. *COX8*, *COX26*, *COA1*, *NDI1*, *QCR6*, *QCR8*, *RCF1* (Dataset1), which is the opposite to what has been described for *A. fumigatus* cells isolated during the caspofungin paradoxical effect (16). Thus, it seems the mechanism responsible for echinocandin tolerance in *C. glabrata* and *A. fumigatus* may be different. This conclusion is also supported by the opposite effects of rotenone in the two systems (16).

In sum, our results reveal a heretofore unappreciated complexity of the mitochondrial involvement in *C. glabrata* response to echinocandins and echinocandin-mediated killing. Whereas we show that the traditionally frequently used mitochondrial inhibitors reduce fungal killing by echinocandins, which is not a desired effect in the clinic, our results suggest that it may be possible to target fungal mitochondria to sensitize *C. glabrata* to these drugs. Indeed, other types of mitochondrial inhibitors, e.g. those that interfere with mitochondrial biogenesis by blocking mitochondrial protein import (56), may be investigated in future studies as potential enhancers of echinocandin fungicidal action.

## MATERIALS AND METHODS

### Yeast strains and media

The *C. glabrata* strains used in this study were ATCC2001 and mutants (gene deletions, petite mutants) derived from ATCC2001 (Table S3). Cells were cultured in standard yeast extract-peptone-dextrose (YPD) medium at 37°C.

### RNA sequencing for echinocandin “enriched” tolerant cells

Overnight cultures containing 10^7^/ml *C. glabrata* cells were exposed to 0.25 µg/ml of caspofungin or 0.06 µg/ml of micafungin. After a 24-hour incubation at 37°C with shaking (150 rpm), cells were then collected by filtration using 0.45-µm mixed cellulose ester membrane filters (Millipore) and resuspended in PBS stained with 10 µg/ml propidium iodide (PI) and sorted using the BD FACSMelody (BD Biosciences). Logarithmically growing “no drug” controls were collected after 2 hours of growth in drug-free YPD, whereas “stationary phase” controls were collected after 24 hours of growth in drug-free YPD. Each sample was collected by FACS, treated with zymolyase (5U per 10 µl of cells) for 30 minutes at 37°C to digest the cell walls, and then pipetted into a 96 well-plate to produce ∼10 PI-negative cells per well. Thirty-six biological replicates were submitted for each echinocandin treated sample (caspofungin and micafungin) and twelve biological replicates were submitted for each control (logarithmic and stationary). The samples were sent to Columbia University Medical Center (New York, NY) for library preparation, and the “enriched” RNA sequencing of ∼10 live cells per well was performed as described previously (57) with some modifications for the reverse transcription reaction using 40U Maxima H-(ThermoFisher), 2X Maxima H-Buffer (ThermoFisher), 4U SuperaseIN (ThermoFisher), 15% polyethylene glycol, and 2uM Template Switching Oligo (IDT) in a total volume of 7.5 µl.

For the RNA-seq analysis, reads were aligned to the NCBI reference, GCF_000002545.3_ASM254v2, using STAR (58). Alignments were assigned to genes using featureCounts (59). The bam files produced by featureCounts were sorted and indexed with samtools (60). The package umi_tools was used to assemble a count matrix (61). Wells were filtered by removing outliers in the distributions of total counts, genes detected, and mitochondrial expression. Highly abundant ribosomal RNAs, which would interfere with normalization, were removed prior to normalization and differential expression analysis. The package sctransform was used to normalize count data (62), and Seurat was then used perform differential expression testing with Wilcoxon rank sum test between groups (63). Raw RNA-seq data have been deposited at the Gene Expression Omnibus (accession no. GSE178797) and differential expression genes (DEGs) files are shown in Dataset 1. The list of *C. glabrata-S. cerevisiae* direct orthologs was downloaded from http://www.candidagenome.org/download/homology/orthologs and supplemented by manual curation of *C. glabrata* genes using http://www.candidagenome.org. Functional categories (FUNCAT) enrichment analysis was performed using FungiFun2 (https://elbe.hki-jena.de/fungifun/) (64). The heatmap was generated using R studio.

### RNA isolation and “bulk” transcriptome analysis

Cultures containing 10^7^/ml *C. glabrata* were exposed to 0.06 µg/ml of micafungin for 24 hours and collected in triplicate in the same fashion as explained in the previous section. Stationary phase cells were collected after 24 hours of growth in drug-free YPD in duplicate as no-drug control. Total RNA was extracted using the RNeasy Mini kit (QIAGEN Science) following the manufacturer’s instructions. The RNA was then treated with RNase-free DNase (ThermoFisher) according to the manufacturer’s recommendations and stored at -80°C until RNA sequencing was performed by Genewiz (South Plainfield, NJ). The RNAseq data were analyzed using Basepair software (Basepair, New York, NY) with a pipeline as follows: reads were aligned to the transcriptome derived from sacCer3 using STAR with default parameters, read counts for each transcript were measured using featureCounts and the differentially expressed genes were determined using DESeq2, and a cutoff of 0.05 for the adjusted P value (corrected for multiple hypotheses testing) was used for creating differentially expressed gene lists (Dataset 1). GSEA was performed on normalized gene expression counts, using gene permutations for calculating P value. The fastq RNAseq files have been deposited at the Gene Expression Omnibus (accession no. GSE178656).

### Antifungal susceptibility testing

Micafungin (Astellas), caspofungin (Merck) and fluconazole (LKT Labs) susceptibility testing was performed using a broth microdilution method following CLSI standards (65) with some modifications. The media used was YPD broth and the concentrations tested ranged from 0.0035 to 2 µg/ml to echinocandins and 0.25-128 µg/ml to fluconazole. Minimum inhibitory concentrations (MICs) were visually read after 24 hours of incubation at 37°C. At least three biological replicates were performed.

### Cell survival measurements after caspofungin treatment combined with inhibitors

To measure the effects of topoisomerase or mitochondrial inhibitors on cell survival in the presence of caspofungin, overnight cultures containing 10^7^ cells/ml were resuspended in fresh YPD and added caspofungin plus one of the inhibitors at the following concentrations. The selected concentrations for the topoisomerase inhibitors were 3 µM, 200 µM and 100 µM of doxorubicin (ThermoFisher), etoposide (Millipore Sigma), and topotecan (Cayman Chemical), respectively, all dissolved in DMSO. The selected concentrations for the mitochondrial inhibitors or ROS scavengers were 0.3 mM, 0.005 mM, 0.5 mM and 50 mM of rotenone, diphenyleneiodonium chloride, sodium azide, and ascorbic acid (Millipore Sigma), respectively, first two dissolved in DMSO whereas the other two dissolved in water. The selected concentration of rotenone was determined by its solubility in liquid YPD because concentrations higher than 0.3 mM resulted in precipitation. After 24 hours of drug exposure, aliquots were harvested and dilutions plated on drug-free YPD plates. Percentage of survival was calculated based on the observed number of colonies from caspofungin cultures relative to the corresponding colony counts from non-caspofungin treated cultures. At least three biological replicates were performed for every strain and condition.

### Reactive oxygen species quantification

The presence of intracellular reactive oxygen species (ROS), more specifically hydrogen peroxide, was assessed by the cell-permeant 2′,7′-dichlorodihydrofluorescein diacetate (CFDA) (10 µg/ml, ThermoFisher) according to the manufacturer’s instructions. Cells were collected and washed with PBS, then 10 minutes before flow cytometric analysis each sample was also stained with 10 µg/ml PI to exclude PI-positive dead cells. The superoxide indicator dihydroethidium (DHE) was used to assess superoxide radical species (6.5 µg/ml in DMSO, Thermo Fisher) following the manufacturer’s instructions. In this case, dead cells to be excluded from the analysis were stained with Sytox Green (1 µM, ThermoFisher). Percentages of ROS-positive cells were calculated relative to all live (PI or Sytox Green-negative) cells after incubation with the dyes. Data were analyzed using FlowJo™ software v10.6.1 (BD Biosciences).

### RNA extraction and quantitative real-time reverse-transcription (RT)-PCR

Overnight cultures containing 10^7^ *C. glabrata* cells/ml were resuspended in fresh YPD containing echinocandins (0.25 µg/ml of caspofungin or 0.06 µg/ml of micafungin) and harvested after 2, 6 or 24 hours, filtered and concentrated in the same fashion as for the “enriched” RNAseq transcriptomic analysis, except that cells were resuspended in RNA extraction buffer provided by the Qiagen kit instead of PBS. Non-treated controls (logarithmic and stationary phase cells) and H_2_O_2_-treated controls (50 mM) were harvested too in the same fashion, whereby the volume of overnight culture needed to obtain a big enough cell pellet to isolate RNA was smaller. Total RNA was extracted using the RNeasy Mini kit (QIAGEN Science) following the manufacturer’s instructions. The RNA was then treated with RNase-free DNase (ThermoFisher) according to the manufacturer’s recommendations. RNA samples were stored at -80°C. The transcript levels of *SOD2*, *CTA1*, *GRX2*, *YAP1*, *MSN2*, and *MSN4* were measured by RT-PCR using One Step SYBR PrimeScript RT–PCR Kit II (TaKaRa). Reactions were run on Mx3005P qPCR System (Agilent Technologies) containing 10 ng RNA sample, 0.4 μM of each primer (Table S4), 12.5 μl 2 × One Step SYBR RT-PCR Buffer, and 1 μl PrimeScript 1 step Enzyme Mix 2 in a volume of 25 μl. Thermal cycling conditions were 42°C for 5 min for the reverse-transcription and PCR cycling with initial denaturation at 95°C for 10 s, followed by 40 cycles of denaturation at 95°C for 5 s and annealing and elongation at 60°C for 20 s; a post PCR melting curve analysis with 95°C for 5 s, 60°C for 1 min then increasing to 95°C.

Each experiment was carried out in biological triplicates and at least two technical triplicates of each biological replicate were also performed. Negative controls were included in each run. The *RDN5.8* gene was used as the reference gene to normalize the data (66). Comparative expression analyses were performed using the 2^−ΔΔCT^ method (67). The fold changes were determined from the normalized expression of the average of treated samples relative to the normalized expression of the average of stationary non-treated controls. Calculations were also done using non-treated logarithmic-phase controls; all trends were similar, so these data were not shown.

### *In vitro* echinocandin tolerance assay

Overnight cultures containing 10^7^ *C. glabrata* cells were resuspended in fresh 1 ml YPD and incubated at 37°C with shaking (150 rpm) for 24 hours in the presence of a range of caspofungin concentrations (0, 0.016, 0.06, 0.25, 1, 4 µg/ml). After 24 hours, 0.1 ml of the appropriate dilutions for each culture was plated onto YPD plates. Colony forming units (CFU) were determined and survival percentage was obtained by normalizing the CFU obtained from cultures treated with the indicated concentration of drug to non-treated controls. Co-treatment with mitochondrial inhibitors or ROS scavengers was performed in the same way as described above, except when there was overnight pre-exposure to the mitochondrial inhibitors before caspofungin was added to the media. At least three biological replicates were performed for every strain and condition.

### Lipid peroxidation

Suspensions of *C. glabrata* 10^7^ stationary-phase cells/ml were exposed to caspofungin (0.25 and 1 µg/ml) or hydrogen peroxide (5 and 10 mM) and incubated with or without 50 μM of diphenyl-1-pyrenylphosphine (DPPP) (ThermoFisher) at 37°C (68, 69). The fluorescence intensity was measured every hour using a Tecan infinite 200 Pro plate reader with a 340/380 nm excitation filter. Data are shown in arbitrary units of fluorescence intensity after 2 and 6 hours of caspofungin exposure.

### Western blotting

*C. glabrata* cells exposed to echinocandin drugs (0.25 µg/ml of caspofungin or 0.06 µg/ml of micafungin), as well as positive (50 mM H_2_O_2_) and untreated (stationary or logarithmic-phase cells) control samples, were collected after 2, 6, and 24 hours of growth, except for H_2_O_2_ where 24-hour cultures had too few cells to proceed. Whole-cell lysates were prepared by trichloroacetic acid (TCA) precipitation. Briefly, cell pellets were resuspended in 20% TCA, broken by bead beating, and washed twice with 5% TCA. Proteins were pelleted, resuspended in sodium dodecyl sulfate-polyacrylamide gel electrophoresis (SDS-PAGE) loading buffer, incubated at 95°C for 5 min, and centrifuged prior to loading on 16% acrylamide gels. Antibodies were obtained commercially: anti-H2A (Active Motif catalog no. 39945) and anti-γH2A.X (Abcam ab15083).

### Spotting assay

Cells obtained from overnight cultures grown in YPD broth at 37°C were spotted in serial 10-fold dilutions on YP plates containing 2% glycerol or YPD plates containing 0.05 mg/ml of Congo red (Millipore Sigma) or 0.01 mg/ml of calcofluor white (Millipore Sigma). Growth of each strain was evaluated after 24 hours of incubation at 37°C. Petite mutants grew much slower than the other mutants, so their growth was also evaluated after 48 hours.

### Microscopy mitochondrial detection

Mitochondria were stained with the fluorescent dye MitoTracker Green FM (ThermoFisher) according to the manufacturer’s protocol but with some modifications for yeast staining. The cells were harvested by centrifugation, diluted to 10^6^ cells/ml in 10mM HEPES buffer, pH 7.4, containing 5% of glucose and stained for 30 min with 0.1 µM MitoTracker Green at room temperature. Stained cells were visualized using a Nikon Eclipse Ti2 inverted microscope (488 nm filter) with Hamamatsu ORCA-Flash4.0 camera and analyzed using NIS-Elements software.

### Construction of *C. glabrata* deletion mutants

Deletion mutants were generated in-house using a CRISPR-Cas9 targeted integration replacing the desired ORF by a nourseothricin (NAT) resistance cassette (Table S3). The deletion constructs were generated by PCR using ∼100 nucleotide-long primers designed to amplify NAT and also containing homology regions flanking the locus of interest. All primers used are listed in Supplementary Table 4. Gene replacement by the deletion cassettes was performed using CRISPR as described previously (70). Briefly, cells were made competent for electroporation using the Frozen-EZ yeast transformation kit (Zymo Research) according to the manufacturer’s instructions and then electroporated with Cas9-gRNA complex (Integrated DNA Technologies) and the DNA containing the deletion construct. Transformants were selected on NAT-containing plates and validated by PCR amplification and sequencing of the targeted locus using external primers (Table S4). At least two independent transformants were generated and analyzed for every deletion mutant. All primers were ordered from Integrated DNA Technologies (Coralville), and all Sanger sequencing of the above-described constructs was done by Genewiz (South Plainfield).

### Measurement of inhibition of glucan synthase by micafungin

Three isogenic ATCC2001 background strains were selected to examine glucan synthase (GS) inhibition by echinocandins *in vitro*: wild-type, *ndi1Δ*, and caspofungin-derived petite mutant #1. The strains were grown with vigorous shaking at 37°C to early stationary phase in YPD broth, and cells were collected by centrifugation. Cell disruption, membrane protein extraction, and partial GS purification by-product entrapment were performed as described previously (36). Sensitivity to micafungin was measured at least in triplicate in a polymerization assay using a 96-well multiscreen high-throughput screen filtration system (Millipore) with a final volume of 100 µl. Serial dilutions of micafungin (0.01–10,000 ng/ml) were used to determine 50% inhibitory concentration (IC50) values. Inhibition profiles and IC50 values were determined using a sigmoidal response (variable-slope) curve-fitting algorithm with GraphPad Prism 8 software.

### Statistical analysis

All shown results represent an average of three or more independent experiments (biological replicates). Error bars represent the standard deviation. All data were analyzed using GraphPad Prism 8 software. Statistical analyses were performed using an unpaired t-test to determine significant differences between experimental groups. P-values ≤0.05 were considered statistically significant.

## ACKNOWLEDGMENTS

This work was supported by NIH 5R01AI109025 to D.S.P. The Flow Cytometry & Cell Sorting Shared Resource was supported by NIH/NCI grant P30-CA051008. This research was funded in part through the NIH/NCI Cancer Center Support Grant P30CA013696 and used the Genomics and High Throughput Screening Shared Resource. It was also supported by the National Center for Advancing Translational Sciences, National Institutes of Health, through Grant Number UL1TR001873. The content is solely the responsibility of the authors and does not necessarily represent the official views of the NIH.

---

**Figure S1. Functional categories enriched among differentially upregulated and downregulated genes.** (A) Functional categories in “bulk” RNAseq samples relative to stationary “no drug” control. (B) Functional categories in tolerant cell-enriched (“enriched”) RNAseq samples relative to stationary “no drug” controls. (C) Functional categories in tolerant cell-enriched (“enriched”) RNAseq samples relative to log phase “no drug” controls. Functional categories found in at least two datasets are marked in violet.

**Figure S2. Mitochondrial defective mutants showed wild type sensitivity to cell wall damaging agents.** Mitochondrial deletion mutants were photographed after 24 hours. Petite mutants of unknown etiology, which were very slow growing, were photographed after 48 hours. YPG = YP + 2% glycerol. Congo red (CR) was used at 0.05 mg/ml and calcofluor white (CW) was used at 0.01 mg/ml.

**Figure S3. Characterization of *C. glabrata* mitochondrial mutants.** (A) MitoTracker Green detection by fluorescence microscopy showed mitochondrial content in petite *C. glabrata* strains. (B) Ethidium bromide-derived petite mutants were hypo-tolerant at sub-MIC caspofungin concentration and showed variable survival at above-MIC caspofungin concentrations. (C) Deletion mutants lacking mitochondrial proteins not directly involved in the respiratory chain showed a caspofungin hypo-tolerant phenotype. The examined deletion mutants were a major ADP/ATP carrier of the mitochondrial inner membrane (*PET9*), a translocase of the inner mitochondrial membrane (*TIM18*), and a catalytic subunit of i-AAA protease complex responsible for degradation of misfolded mitochondrial gene products encoding (*YME1*).

**Figure S4. Diphenyleneiodonium chloride (DPI) and sodium azide did not improved survival in deletion mutants lacking mitochondrial respiratory chain components.** Deletion mutants of mitochondrial respiratory chain components *NDI1* (NADH dehydrogenase of equivalent of complex I) and *COX4* (cytochrome-c oxidase of complex IV) (A), and complex V ATP synthase (*ATP1*, *ATP2* and *ATP10*) (B) were not rescued by DPI in the caspofungin tolerance assay. (C) Sodium azide did not rescue survival of the *cox4Δ* mutant.

**Figure S5.** Exposure of *C. glabrata* cells to the mitochondrial inhibitors (A) rotenone and (B) diphenyleneiodonium chloride (DPI) prior to caspofungin treatment reduced cell survival. Rotenone was still able to rescue growth at several caspofungin concentrations above the MIC.

**Table S1. Sorting cells that did not stain with propidium iodide resulted in the strongest enrichment for cells capable of producing colonies.** Fluorescence-activated cell sorting (FACS) was used with different combinations of live/dead dyes and fluorescent markers in cells cultured in the presence of either 0.25 µg/ml caspofungin or 0.06 µg/ml micafungin for 24 hours, followed by plating of the sorted fractions on drug-free medium and counting the colony forming units (CFU). PI (propidium iodide), CFDA-AM (5-Carboxyfluorescein Diacetate, Acetoxymethyl Ester), SG (Sytox Green), RFP (red fluorescence protein), GFP (green fluorescence protein).

**Table S2. MIC susceptibility testing of selected concentration of inhibitors in combination to caspofungin and micafungin showed identical MIC-values than echinocandins by themselves.** The inhibitor concentration selected was based on the MIC-value obtained in the susceptibility testing together with its solubility in YPD liquid media. DPI (diphenyleneiodonium chloride). *Rotenone is non-soluble >0.3 mM. **Other authors have used 2.5-10 mM of ascorbic acid combined with other drug compounds in *Candida* species (1–3).

**Table S3.** List of *Candida glabrata* strains used in this study and their fluconazole susceptibilities. ECH (echinocandins).

**Table S4.** Sequences of primers used in this study for RTqPCR transcriptional experiments and generation of CRISPR knock-out mutants. *Primers described in a different study (4).

**Dataset 1.** “Bulk” and “enriched” RNAseq data.

